# Nanopore base-calling from a perspective of instance segmentation

**DOI:** 10.1101/694919

**Authors:** Yao-zhong Zhang, Arda Akdemir, Georg Tremmel, Seiya Imoto, Satoru Miyano, Tetsuo Shibuya, Rui Yamaguchi

## Abstract

**Background:** Nanopore sequencing is a rapidly developing third-generation sequencing technology, which can generate long nucleotide reads of molecules within a portable device in real time. Through detecting the change of ion currency signals during a DNA/RNA fragment’s pass through a nanopore, genotypes are determined. Currently, the accuracy of nanopore base-calling has a higher error rate than short-read base-calling. Through utilizing deep neural networks, the-state-of-the art nanopore base-callers achieve base-calling accuracy in a range from 85% to 95%.

**Result:** In this work, we proposed a novel base-calling approach from a perspective of instance segmentation. Different from the previous sequence labeling approaches, we formulated the base-calling problem as a multi-label segmentation task. Meanwhile, we proposed a refined U-net model which we call UR-net that can model sequential dependencies for a one-dimensional segmentation task. The experiment results show that the proposed base-caller URnano achieves competitive results compared to recently proposed CTC-featured base-caller Chiron, on the same amount of training and test data for in-domain evaluation. Our results show that formulating the base-calling problem as a one-dimensional segmentation task is a promising approach.

**Availability:** The source code and data are available at https://github.com/yaozhong/URnano

**Supplementary information:** Supplementary data are available at attachment online.

## 1. Background

Nanopore sequencing, a third-generation sequencing technique, has achieved impressive improvement in the past several years [1, 2]. A nanopore sequencer measures currency changes during the transit of a DNA or an RNA molecule through a nanoscopic pore and can be equipped in a portable size. For example, MinION is such a commercially available device produced by Oxford Nanopore Technologies (ONT). One key merit of nanopore sequencing is its ability to generate long reads on the order of tens of thousands of nucleotides. Besides the sequencing application, it is actively used in more and more fields, such as microbiology and agriculture.

Base-calling is usually the initial step to analyze nanopore sequencing signals. A base-caller translates raw signals (referred to as squiggle) into nucleotide sequences and feeds the nucleotide sequences to downstream analysis. It is not a trivial task, as the currency signals are highly complex and have long dependencies. ONT provides established packages, such as Scrappie and Guppy. Currently, nanopore base-calling still has a higher error rate when compared with short-read sequencing. Its error rate ranges from 5% to 15%, while the Illumina Hiseq platform has an error rate of around 1%. More and more work is now focusing on solving challenges to further improve base-calling accuracy.

Early-stage base-callers require first splitting raw signals into event segments and predict k-mer (including blanks) for each event. Sequential labeling models, such as hidden Markov model (HMM) [3] and recurrent neural network (RNN) [4] are used for modeling label dependencies and predicting nucleotide labels. It is widely considered that a two-stage pipeline usually brings about an error propagation issue that wrong segments affect the accuracy of base-calling. Recently, end-to-end deep learning models are used to avoid pre-segmentation of raw signals, which enables base-callers to directly process raw signals. For example, BasecRAWller [5] puts the event segmentation step in a later stage after initial feature extraction by a RNN. Chiron [6] and recent ONT base-callers use a Connectionist Temporal Classification (CTC) module to avoid explicitly segmentation for base-calling from raw signals. With CTC, a variant length base sequence can be generated for a fixed-length signal window through output-space searching.

On the other hand, even though those base-callers can translate raw signals to bases directly, segmentation and explicit correspondence between squiggles and nucleotide bases are also informative. It can provide information for detecting signal patterns of target events, such as DNA modifications [7]. In a re-squiggle algorithm, base-calling and event detection are also required.

In this paper, we performed base-calling from the point of view of instance segmentation and developed a new base-caller named URnano. Distinguished from previous work that treats base-calling as a sequence labeling task, we formalize it as a multi-label segmentation task that splits raw signals and assigns corresponding labels. Meanwhile, we avoid making the assumption that each segment is associated with a k-mer (*k* ≥ 2) and directly assign nucleotide masks for each currency sampling point. On model-level, based on the basic U-net model [8], we proposed an enhanced model called UR-net that is capable of modeling sequential dependencies for a one-dimensional (1D) segmentation task. Our base-caller is also an end-to-end model that can directly process raw signals. Our experiment results show that the proposed URnano achieves competitive results when compared with current base-callers using CTC decoding on the same dataset, while retains speed advantage with its model architecture.

## 2. Methods

The overall structure of URnano is described in Figure 1. URnano contains two major components: ➀ UR-net for signal segmentation and base-calling.➁ Post-processing. For streaming signals generated by a nanopore sequencer, URnano scans signals in a fixed window length *L* (e.g., *L* = 300) and slides consequently with a step length *s* (e.g., *s* = 30). Given signal input *X* = (*x*_1_, *x*_2_, *…, x*_*i*_, *…, x_L_*), UR-net predicts segment label masks *y*_*i*_ for each *x*_*i*_. The output of UR-net *Y* = (*y*_1_, *y*_2_, *…, y*_*i*_, *…y_L_*) has exactly the same length as the input *X* and *y*_*i*_ ∈ {*A*_1_, *A*_2_, *C*_1_, *C*_2_, *G*_1_, *G*_2_, *T*_1_, *T*_2_}. Here, {*A*_1_, *C*_1_, *G*_1_, *T*_1_} and {*A*_2_, *C*_2_, *G*_2_, *T*_2_} are alias label names, which is designed to process homopolymer repeats (described in section 2.3). After label mask *Y* is generated, we conduct a post-processing step that transforms *Y* to *Y* ^′^ ∈ {*A, C, G, T*}*^N^*, where *N* is the length of the final basecall. The post-processing contains two simple steps. First, it collapses consecutive identical label masks as one label. Second, the collapsed labels in alias namespace are transformed back to bases in {*A, C, G, T*}. *Y* ^′^ is the final base-calls of the URnano.

**Fig. 1:**
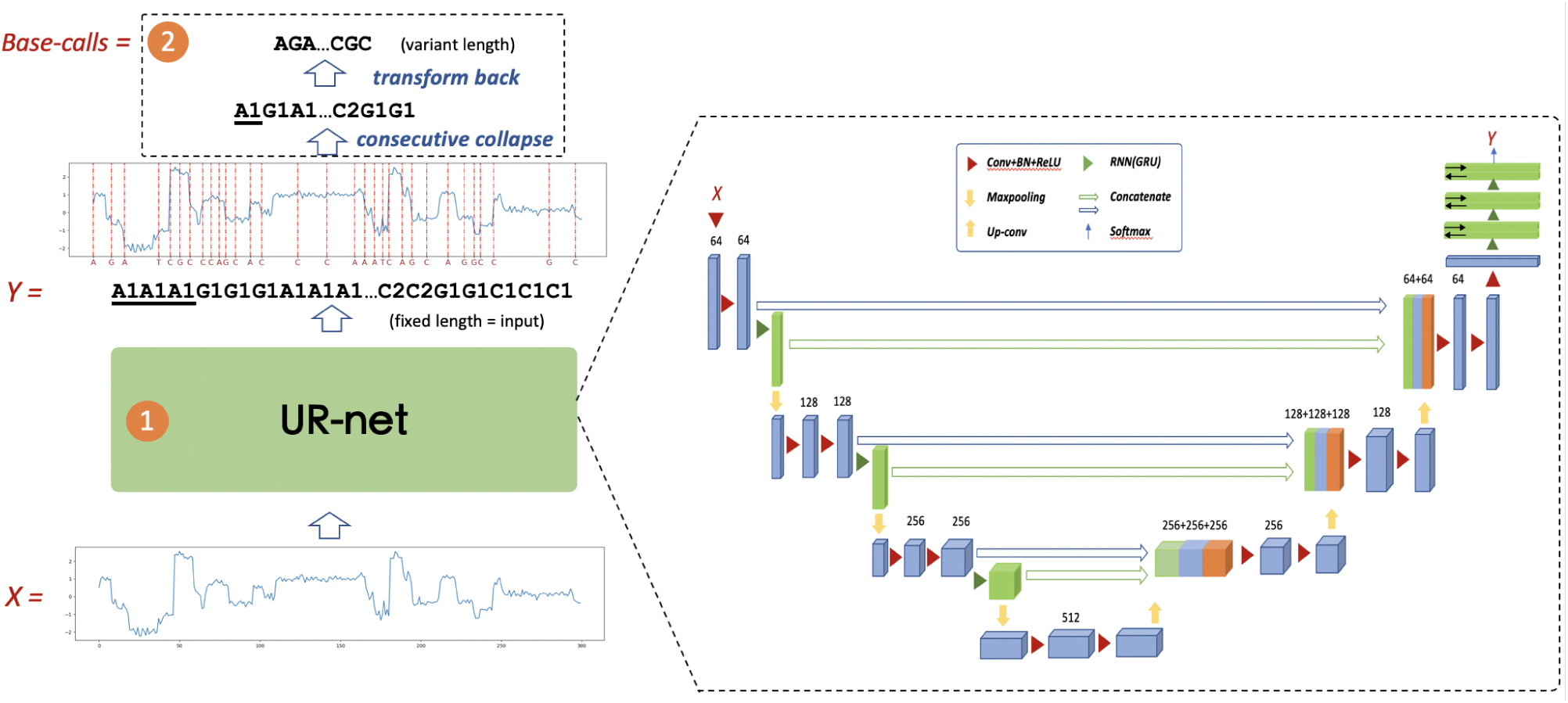
Overall structure of URnano baser-caller. Block➀is the UR-net deep neural network structure. Block ➁ is the post-processing part that transforms the UR-net’s output to final base-calls.

Besides predicting base-calls, URnano also generates a signal segment for each base. In previous work [5, 4], signal segments are assumed to be associated with k-mers of a fixed k (e.g., k=2,4,5). Every base is read as a part of k consecutive events. In URnano, we avoid making the k-mer assumption and directly assign label masks for signals.

### 2.1 UR-net: enhanced U-net model for 1D sequence segmentation

The key component of the URnano is UR-net and its network structure is shown in Figure 1. **In general**, **UR-net is based on the U-net model [8] and is enhanced to model sequential dependencies**^1^. The original U-net is designed for image data in two dimensional (2D) and has achieved the-state-of-the-art performances in many image segmentation tasks. Although the model can be directly applied for 1D data, the 1D segmentation task has its own characteristics that are distinguished from the 2D image segmentation task. In a sequence segmentation task, one segment may not only relate with its adjacent segments but also depends on non-adjacent segments that are long-distance away. Such dependencies were not considered in the original U-net model, which mainly focuses on detecting object regions and boundaries.

The UR-net has a similar U-shape structure as U-net, in which left-U side encodes inputs *X* through convolution and max pooling, and right-U side decodes through up-sampling or de-convolution. We make two major enhancements in the UR-net model, which are highlighted in the green color shown in Figure 1 and described as follows:

- For the left-U side encoding part, we add an RNN layer right after each CONV-BN-RELU block to model sequential dependencies of hidden variables in different hierarchical levels. Those RNN layers are then concatenated with CONV and UP-Sample layer in the right-U side decoding part.
- We add three bi-directional RNN layers in final layers.

Those changes are motivated to enhance the sequential modeling ability for the U-net.

### 2.2 Model training

Given *D* = {(*X*_*i*_, *Y*_*i*_)|*i* = 1*…n*}, we train UR-net with an interpolated loss function that combines dice loss and categorical entropy loss. The task loss of edit distance can not be directly optimized. For each sample, the dice loss (DL) and categorical entropy loss (CE) are defined as follows:

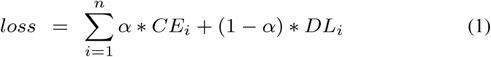

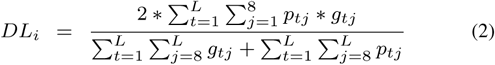

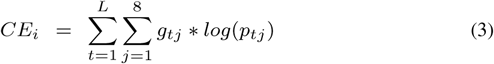

, where *p*_*ij*_ is the *j*-th softmax value for the *i*-th sample. We use Adam [9] to optimize the above loss function.

### 2.3 Homopolymer repeats processing

In genomes, homopolymer repeats (e.g., AAA and TTTT) exists. Figure 2 demonstrates a histogram of homopolymer repeats ^2^ on randomly sampled 200 E. coli reads and 200 *λ*-phage reads. From the figure, we can observe that majority homopolymer repeats have lengths less than 5 base-pairs. For the original U-net model, adjacent bases in a homoploymer can not be distinguished and are merged as one base. This brings about deletion errors if training model directly. To solve this problem, we use an alias trick to differentiate adjacent identical labels. For example, homopolymer repeat “*AAAAA*” in the training data is converted to “*A*_1_*A*_2_*A*_1_*A*_2_*A*_1_” for training UR-net model. In the inference stage, those new labels are transformed into the original representation through post-processing.

**Fig. 2:**
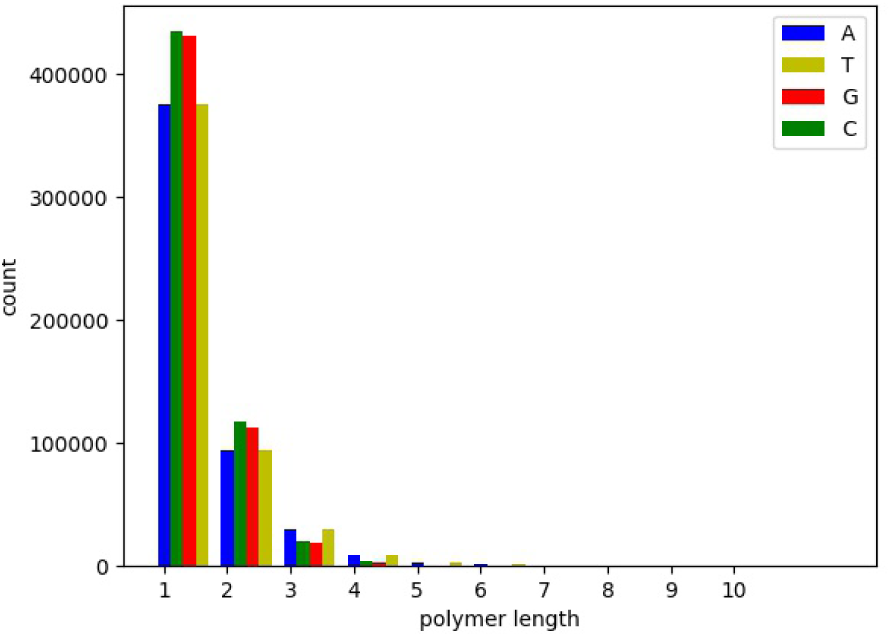
Histogram of homopolymer repeats from 400 E. coli and *λ*-phage reads.

### 2.4 Merge base-calls in sliding window into a whole read

In the training phase, a read is split into the non-overlapping windows of fixed length. In the testing phase, for calculating read accuracy, read signals are scanned with overlapping windows. The sliding window takes a small step *s* (*s < L*). For each time step *t, x*_*t*_ and *x*_*t*−1_ have *L* − *s* overlaps on the signal content. Thus, for a read signal of length *N* we have 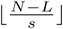 windows and each overlap with its neighbors by *L* − *s*. The base-calls for each input at neighboring positions are merged in pair-wise fashion consequently. We find the start index of the longest consecutive overlap between subsequent nucleotide sequences, and concatenate two predictions from that starting point. After doing the concatenation for all neighboring segments, we count the number of the occurrence of each nucleotide for each position and use the most frequent ones as the final prediction. An example is shown in Figure 3. In the training, the start position of each nucleotide in the signal is known. But in streaming decoding mode, signals near start or end position of the window may be incomplete signals for one nucleotide. Therefore, we clip the first and last base calls of each window before they are used in merging a whole read.

**Fig. 3:**
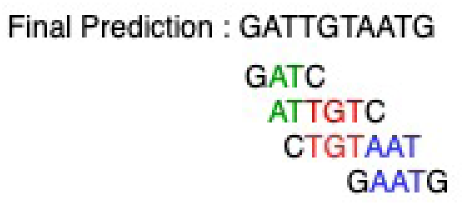
An example on merging base-calls of overlapped slide window for a whole read.

## 3. Experiments

### 3.1 Experiment settings

#### 3.1.1 Data

We compared URnano with recently proposed base-callers: Chiron^3^ and ONT base-callers, such as Scrappie (raw) and Guppy. For comparing model performances, we used publicly accessible curated data set provided by Teng et al. [6]. The training set contains a mixture of randomly selected 2000 E. coli reads and 2000 *λ*-phage reads generated using nanopore’s 1D protocol on R9.4 flowcells. The test set contains the same amount of reads from E. coli and *λ*-phage. To assess read accuracy across species, we use 1000 randomly selected reads from Chromosome 11 of human benchmark sample NA12878 (1D protocol on R9.4 flowcells)^4^.

The input signals are normalized using median shift and median absolute deviation scale parameters 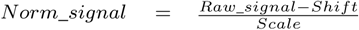. The *Norm*_*signal* usually has values in a range of [−2, 2]. For training deep learning models, signals of a read are split into non-overlapping window segments of a fixed length *L* (*L* = 300 by default). For those samples containing *Norm*_*signal* larger than 10, we filtered them out for training. In total, we have 830,796 training samples. For evaluating read accuracy of a whole read, a sliding window takes a step of 30 for generating overlapped base-calls.

#### 3.1.2 Evaluation metric

We evaluated a base-caller’s performance according to the following three metrics:

- Normalized edit distance (NED) between gold nucleotides and base-calls in non-overlapping windows
- Read accuracy (RA) evaluates the difference between a whole read and its reference.

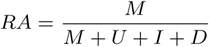

Where *M* is the number of bases identical to reference. *U*, *I* and *D* are the number of mismatches, inserts and deletions, respectively, according to the reference read. Following the evaluation scheme in Chiron, we used GraphMap [10] to align base-calls of a read to the reference genome. The error rates of the aligned reads are calculated using the publicly available Japsa tool^5^.
- Assembly identity and relative length. We assembled genomes using the base-calls from each basecaller. The details of the assembling process are given in the ‘Read assembly results’ section.

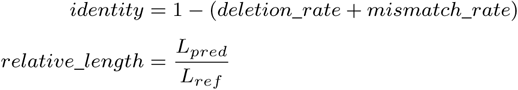

where *L*_*pred*_ is the length of the assembled base-call and *L*_*ref*_ is the length of the reference genome.

#### 3.1.3 Model and base-caller settings

The URnano is implemented using Keras (v2.2.4) with Tensorflow backend (v 1.8.0). The source code is available on github^6^. Chiron (v0.3 clean branch) is evaluated using their provided model. The model is trained on the same data set as URnano. The Beam search version of Chiron is used here, with a beam size of 50. ONT base-callers (Scrappie and Guppy) are used with the provided models. Those models are trained on a large data set, which contains more species, such as H.sapiens.

### 3.2 Base-calling results on non-overlapping segments

Different base-callers and variant deep network architectures are first evaluated using normalized edit distance (NED). In total, 847,201 samples of 300-length window are evaluated. In general, the lower NED is, the more accurate a base-caller is.

**Table 1.** NED of different base-callers and network architectures for non-overlapping window in the test set.

| Category | Model | Mean | Std |
| --- | --- | --- | --- |
| with CTC | Chiron | 0.2084 | **0.1247** |
| no CTC | Unet | 0.3528 | 0.2448 |
|  | 3GRU | 0.2808 | 0.1631 |
|  | Unet+3GRU | 0.1800 | 0.1296 |
|  | URnano | **0.1665** | 0.1329 |

Based on whether a model using Connectivist Temporal Classification (CTC), we categorized base-callers into two types: with CTC and without CTC. Chiron is the first published work using CTC decoding for raw signals. In our segmentation-based models, CTC is not used. The URnano performs better than Chiron with a relative 20% smaller NED. This shows the potential of modeling base-calling as an instance segmentation task. For the NED variance, URnano has a higher value of 0.1329, when compared with 0.1247 of Chiron.

For base-callers without CTC, we compared the performance of different neural network architectures. The original U-net performs the worst of 0.3528, while URnano achieves the best of 0.1665. As the sequential dependencies are not modeled in the U-net, these results indicate the importance of sequential information in the 1D segmentation task.

To take into account the sequential dependencies, we initially added 3 layers of bi-directional gated recurrent units (GRU) for the output of the U-net. This gives about 0.1728 absolute reduction on the NED compared with the U-net. Meanwhile, we observed that the Unet+3GRU performs significantly better than only using 3GRU (0.1 absolute NED reduction). In addition, we incorporated GRU layers in different hierarchical levels of convolutional layers. It gives a further 7.5% relative reduction of NED, when comparing URnano with Unet+3GRU.

### 3.3 Segmentation results

In this section, we investigated event segments for each predicted nucleotide. Figure 4 demonstrates an example of base-calling and segmentation by URnano and Chiron. For URnano, the signal segment for each base can be directly derived through label masks. As in the post-process of URnano, consecutive identical masks are merged as one base, a region of consecutive identical masks is just an event segment. For Chiron, it is a bit tricky to generate event segments from predictions, as the model does not explicitly provide segmentation information. In order to compare our results with the Chiron, we use a heuristic approach to generate segments based on Chiron’s results. Based on the base-calling prediction, segmentation points can be found by observing the softmax probabilities fed to the CTC decoder. From the start, we find the next significant peak for the current nucleotide as the segmentation point and repeat this for the rest. A significant peak in the probabilities is determined by a threshold (0.2 in our case). For example, as shown in the Chiron prediction in Figure 4, after mapping ‘T’ to the first peak (shown in green), this algorithm searches for the next significant peak for ‘G’ (shown in blue). The search stops after reaching a significant peak and we move to the next iteration.

**Fig. 4:**
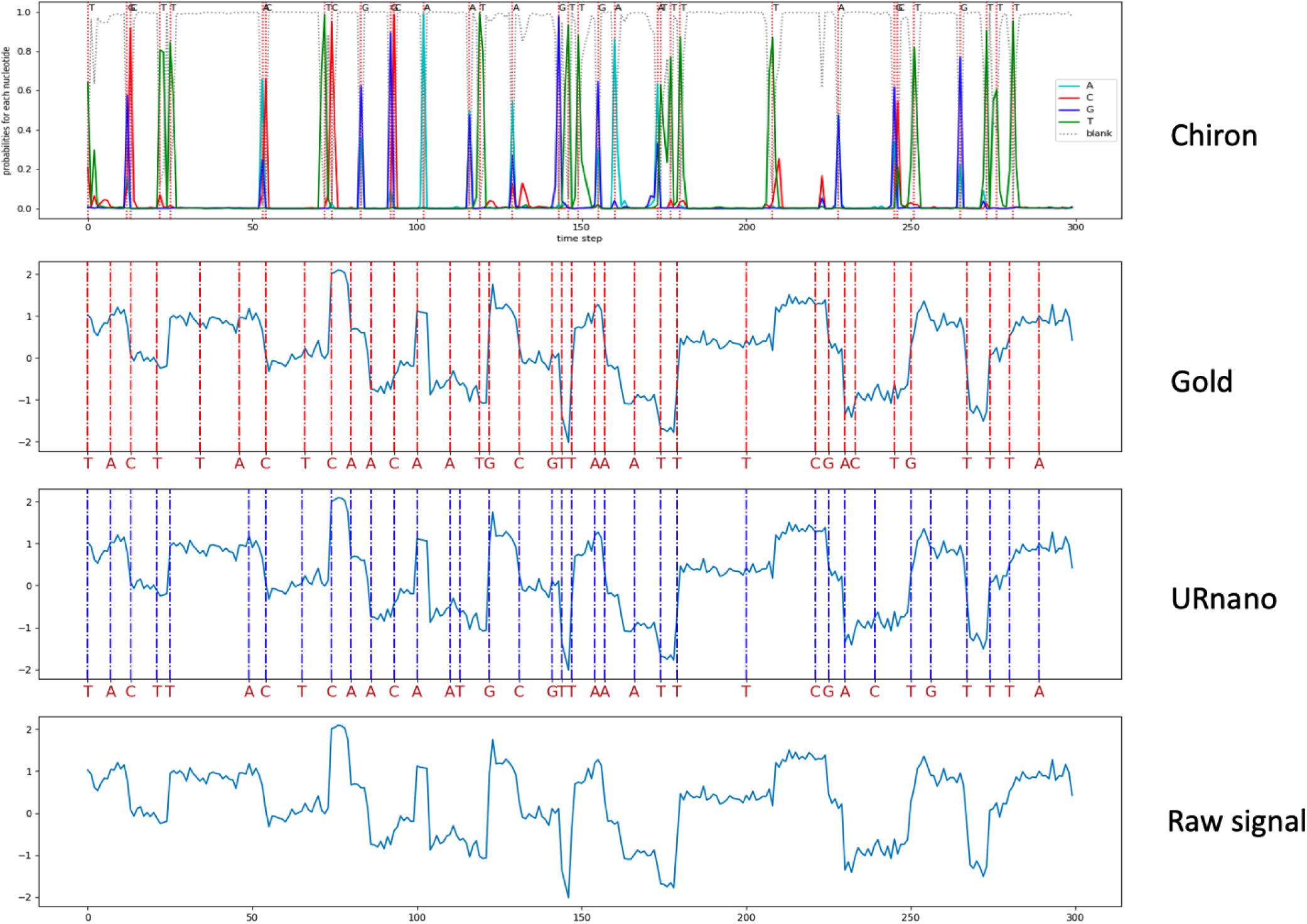
An segmentation example An example of base-calling results by Chiron and URnano for the sequence of “TACTTACTCAACAATGCGTTAAATTTCGACTGTTTA”. A dotted vertical line indicates the start position of a nucleotide segment

**Table 2.** Results of read accuracy on the test set. Note that ONT base-callers in gray-color line is trained on a larger dateset, which are used as references. We mainly compared the performances of Chiron and URnano.

| Species | Base-caller | Deletion | Insertion | Mismatch | Identity | unaligned | Read Accuracy |
| --- | --- | --- | --- | --- | --- | --- | --- |
| E. coli | Chiron | 0.0725 | 0.0304 | 0.0554 | 0.8721 | 76/2000 | 0.8417 |
|  | URnano | 0.0585 | 0.0457 | 0.0452 | 0.8963 | 10/2000 | 0.8506 |
|  | Scrappie | 0.0448 | 0.0343 | 0.0352 | 0.9200 | 8/2000 | 0.8857 |
|  | Guppy | 0.0459 | 0.0192 | 0.0270 | 0.9271 | 4/2000 | 0.9079 |
| λ-phage | Chiron | 0.0841 | 0.0333 | 0.0586 | 0.8573 | 303/2000 | 0.824 |
|  | URnano | 0.0662 | 0.0365 | 0.0395 | 0.8943 | 11/2000 | 0.8578 |
|  | Scrappie | 0.0503 | 0.0391 | 0.0394 | 0.9102 | 7/2000 | 0.8712 |
|  | Guppy | 0.0516 | 0.0247 | 0.0327 | 0.9156 | 3/2000 | 0.891 |
| Species | Base-caller | Deletion | Insertion | Mismatch | Identity | unaligned | Read Accuracy |
| Human | Chiron | 0.0903 | 0.0587 | 0.0663 | 0.8434 | 422/1000 | 0.7847 |
|  | URnano | 0.1055 | 0.0650 | 0.0776 | 0.8169 | 380/999 | 0.7519 |
|  | Scrappie | 0.0543 | 0.0703 | 0.0513 | 0.8943 | 349/1000 | 0.8241 |
|  | Guppy | 0.0671 | 0.0499 | 0.0549 | 0.8780 | 292/973 | 0.8281 |

Figure 4 demonstrates the segmentation results generated by URnano for a randomly selected input. From the gold segmentation, we can observe **the signals for a nucleotide is not evenly distributed across time**. This is mainly due to the fact that, the speed at which a molecule passes through a pore changes over time. The speed issue makes the segmentation a non-trivial task. Traditional statistical approaches without considering the speed changes may not work. Here, the proposed URnano is designed to learn segmentation from the data, which implicitly considers the speed changes embedded in signals. For example, events of ‘T’s around 150 time-step tend to have short lengths than that in 200 time-step. The URnano can distinguish such speed changes as shown in the third row of the figure. For Chiron, the speed changes can also be detected as shown in the Figure, yet the boundaries are not as accurate as segmentation oriented URnano in this example.

For the beginning part of the signal in this example, both Chiron and URnano generate correct base predictions, but the segments of ‘TT’ shift a bit compared to the gold standard. In general, URnano provides a more intuitive and easier interpretable segmentation result when compared to Chiron.

### 3.4 Base-calling results on read accuracy

We evaluated read accuracy for the whole reads on the test set. For in-domain evaluation, we tested on 2000 E. coli and 2000 *λ*-phage reads, separately. In both species, URnano has a lower Deletion and Mismatch rate, when compared with Chiron. But Insertion rate of URnano is 0.0153 higher in E. coli and 0.0032 in *λ*-phage than corresponding values in Chiron. Overall, URnano achieves better identity score and read accuracy in those two species than Chiron. Although it is not fair to directly compare with ONT base-callers trained on a larger data set, performances of Scrappie and Guppy here are used as relative references for Chiron and URnano results. For cross-species, we tested on 1000 randomly selected reads from chromosome 11 of human benchmark sample NA12878. Chiron performs better than URnano on identity and read accuracy, but has more reads that are not aligned.

### 3.5 Read assembly results

We also evaluated the quality of the assembled genomes using the reads generated by each base-caller on the test set. We make use of the same evaluation pipeline of Teng et al. [6] in order to produce consistent results. We used minimap and miniasm [11] tools to first map the reads against each other to find raw contigs and further polished them for 10 rounds by using Racon [10]. Then the polished contigs are shredded into 10K long read fragments and each fragment is aligned to the reference genome and we calculated the identity rates and the relative lengths. The mean identity rate is calculated by taking the average of the identity rates for each contig aligned to the reference. The identity rate for a single contig is the average of the identity rates of each aligned part. If the total length of the aligned parts is smaller than half of the read length we assume it to be unaligned and the identity rate for that contig is 0. The mean relative length is also calculated in a similar way.

We used the E. coli, *λ*-phage and Human test set to evaluate the quality of the contigs assembled. Table 3 gives the results for assembly quality comparison. On all the test sets, URnano outperforms Chiron for both identity rate and relative length. Also the gap between URnano and ONT basecallers is reduced when compared to that in the raw read accuracy task. Although the read accuracy results of Chiron for human data is better compared to URnano, the higher number of unaligned reads (shown in Figure 2) harms the assembly results.

**Table 3.**
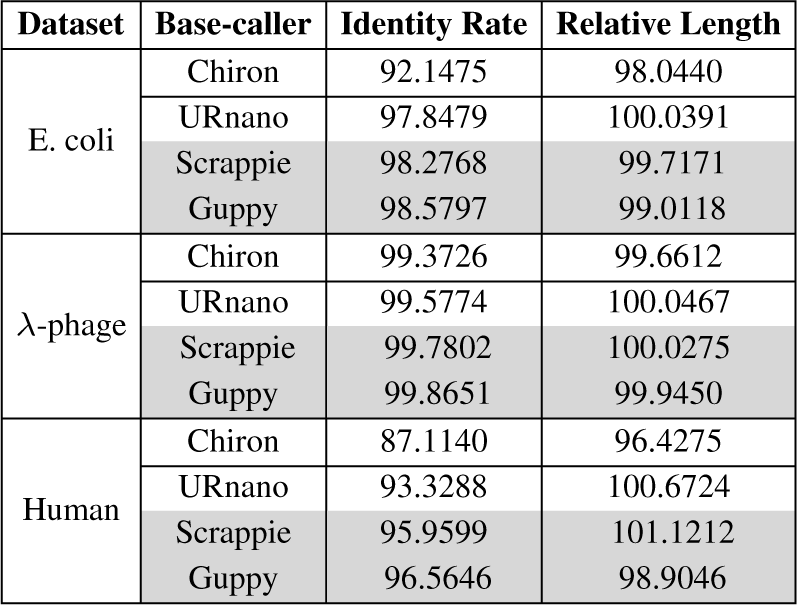
Evaluation with assembly on the test set. Note that ONT base-callers in gray-color line is trained on a larger dateset, which are used as references.

As for the time issue, URnano does not contain CTC decoding step, which makes it around 2x faster on average over all test data (Avg. 1165 bp/s) than Chiron (Avg. 590 bp/s) using Nvidia Tesla V100. We give the average over all input signals as it is a more robust metric compared to giving the maximum speed.

## 4. Discussion

To compare with related deep-learning base-callers, we analyzed 5 base-callers and enumerated their key modules including network input, network structure, network output and post-process of each one, shown in Table 4. For all the 5 base-callers except DeepNano, raw signals can be directly processed. DeepNano requires raw signals to be segmented into events before base-calling.

**Table 4.**
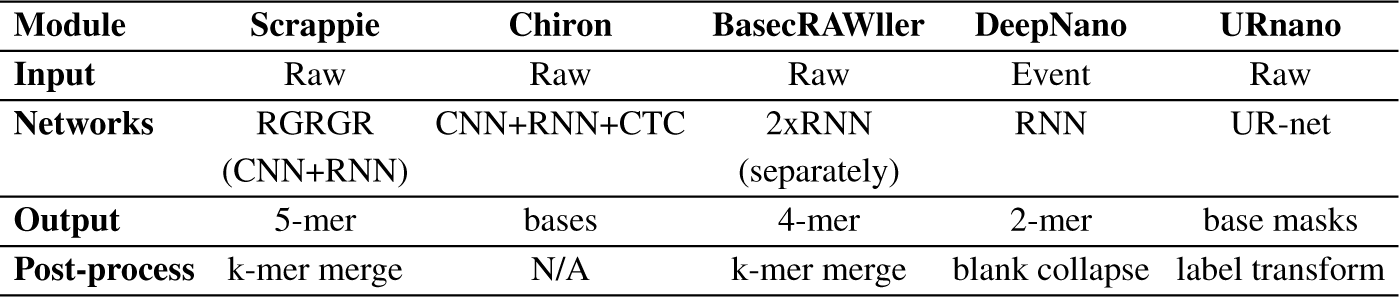
A brief summary of related deep-learning base-callers

For neural network architectures, convolutional layer (CNN) and recurrent layer (RNN) are commonly used modules. CNN is generally used to extract features from raw signals and prepares input for RNN. RNN module is used to learn dependencies among hidden units. In URnano, our experiment also demonstrates the usefulness of using RNN for 1D segment mask prediction. Besides using RNN in final layers, URnano also demonstrates the combination of CNN and RNN layers in the encoding stage can further improve the base-calling performance.

For the output of neural networks, Scrappie, DeepNano and BasecRAWller predict k-mers (*k* ≥ 2). Therefore, a k-mer merging step requires post-processing to generate final base-calls for ONT Scrappie and BasecRAWller. For DeepNano that predicts 2-mer with blanks, a blank collapse step is applied instead. Chiron uses CTC decoding to generate base-calls of variant length through beam searching hidden unit spaces. The output of Chiron also includes blank labels, which are collapsed in the CTC decoding stage.

In a real physical process, the speed of a molecule passing through a nanopore changes over time. This can be observed in the Figure 4. A k-mer assumption using fixed k may not hold over time. Although incorporating blank labels can deal with low-speed case, the high-speed one that involves more bases for the same signal length could exceed the limit of the fixed k. For Chiron and URnano, the fixed k-mer assumption is avoided in base-calling. Chiron uses CTC decoding, while URnano uses label masks that are smaller units than 1-mer.

To curate the data for training a base-caller, a re-squiggle algorithm is usually applied. In a re-squiggle algorithm, raw signal and associated base-calls are refined through alignment to a reference. After re-squiggling, a new assignment from squiggle to a reference sequence is defined. In the previous re-squiggle algorithms, such as Tombo^7^, event detection and sequence to signal assignment are performed in sequence, separately. We think the proposed URnano can be used as the base-caller in a re-squiggle algorithm, as it can do base-calling, event detection and sequence to signal assignment jointly in an end-to-end manner.

In the read accuracy evaluation, URnano performs worse than Chiron in the cross-species human data, but works better in both in-domain species. This indicates URnano may be over-trained for the data in the species. This is also reflected in the training parameters that we trained 100 epochs, while Chiron was trained in 3 epochs. To solve this issue, on one hand, we can incorporate more regularization modules in the network. On the other hand, it would be more effective to incorporate more training data covering various species. We intend to further refine URnano as future work.

## 5. Conclusion

In this paper, we proposed a novel base-calling approach from the perspective of instance segmentation. We formalized base-calling as a multi-label segmentation task and developed an end-to-end solution that can perform base-calling from raw signals and generate signal segments for base-calls at the same time. In addition, we proposed an enhanced deep neural network architecture called UR-net for 1D sequence data. The proposed URnano achieves better performances in normalized edit distance, read accuracy and identity when compared with Chiron on in-domain data. For read assembly on the same test set of low depth coverage, URnano outperforms Chiron for both identity rate and relative length. Without using CTC, the decoding speed of the URnano achieves about two times faster than Chiron.

## Footnotes

1 “R” represents a refinement of U-net and the integration of RNN modules.

2 non-repeats are treated as one time repeat for reference

3 https://github.com/haotianteng/Chiron

4 https://github.com/nanopore-wgs-consortium/NA12878

5 https://japsa.readthedocs.io/en/latest/license.html

6 https://github.com/yaozhong/URnano

7 https://github.com/nanoporetech/tombo

